# Protein Language Modeling beyond static folds reveals sequence-encoded flexibility

**DOI:** 10.64898/2026.01.21.700698

**Authors:** Finn H. Lüth, Victor Mihaila, Milot Mirdita, Martin Steinegger, Burkhard Rost, Michael Heinzinger

## Abstract

**Motivation:** Proteins function through motion. Yet, most discoveries still commence with static representations of protein structures. Here, we investigated the feasibility of leveraging protein dynamics to improve homology detection.

**Results:** We introduce ProtProfileMD, a sequence-to-3D-probability model that predicts, from an amino acid sequence, a profile of discrete structural representations capturing protein dynamics. We applied supervised parameter-efficient finetuning of the ProstT5 protein Language Model (pLM) to predict per-residue distributions over Foldseek’s 3Di alphabet derived from motions observed in molecular dynamics. This original result reveals that the 3Di tokens, despite being coarse-grained descriptors of 3D structure, still offer sufficient resolution to capture aspects of conformational changes. This is evidenced by a correlation between fluctuations in the 3D protein structure over the course of a molecular dynamics trajectory and the entropy of 3Di states. Based on this insight, we introduce a proof-of-concept for making remote homology detection of proteins more sensitive by leveraging a protein’s distinctive dynamic fingerprint captured by our model. Our method recovers flexibility signals with a fidelity that is biologically relevant, improving search and complementing protein structure predictions, for example, by flagging flexible, disordered, or other functionally relevant regions.

**Availability and Implementation:** ProtProfileMD is available at github.com/finnlueth/ProtProfileMD. The associated training data and model weights are available at huggingface.co/datasets/finnlueth/ProtProfileMD and huggingface.co/finnlueth/ProtProfileMD.

## Introduction

The fundamental principle *sequence determines structure, structure determines function* has guided molecular biology for decades [1, 2, 3]. However, this assertion omits a key detail: proteins act as dynamic machines. They do not exist as rigid structures; instead, they explore ensembles of conformations across many orders of timescales [4, 5, 6, 7]. These conformational changes are often vital for a protein’s function, ranging from enzymatic catalysis and allosteric regulation to molecular recognition and signaling [8, 9, 7]. X-ray crystallography yields high-resolution structures but often obscures the dynamics, whereas Nuclear Magnetic Resonance spectroscopy (NMR) often captures the dynamics, albeit at the cost of lower resolution and reduced protein size [10]. *In silico* alternatives such as Molecular Dynamics (MD) simulations, integrating Newton’s equation of motion, offer fine-grained atomistic insight into protein motions but are computationally expensive, thereby limiting timescales and system sizes [10]. More recently, Deep Learning (DL) based generative ensemble methods [11, 12, 13, 14, 15] provide promising alternatives but remain limited, possibly due to insufficient high-quality training data and computational resources. Consequently, these methods scale poorly to proteome-wide analyses.

Protein Language Models (pLMs) learn information from very large, unlabeled protein sequence databases. By implicitly learning aspects of “the language of life”, they capture a variety of protein features spanning evolutionary, biophysical, structural, and functional constraints as latent representations [16, 17, 18, 19, 20, 21]. Prediction methods using exclusively input from pLM embeddings achieve state-of-the-art performance across a variety of tasks, ranging from signal peptide prediction to antibody design, as well as inference of dynamics [22, 23, 24, 25]. With the help of sequence-based alignment and search tools [26, 27, 28], ProstT5 has been adapted to large-scale proteomic analysis [29]. This is achieved through discrete, one-dimensional (1D) structural representations that can be inferred from amino acid sequence, omitting the costly transitive step of explicitly predicting protein structures on the atomic level [29, 27].

Expanding the rationale behind ProstT5, here we introduce **ProtProfileMD**, a novel composition of MD and pLMs which enables large-scale, proteome-wide, dynamics-informed comparison and analysis using amino acid sequence as the only input. This is realized by linking evolutionary signals to conformational flexibility via the transfer of MD-derived priors into a pretrained pLM. Protein dynamics are represented by probability distributions imputed from MD trajectory frames, each of which has been encoded by the discrete structural alphabet, 3Di, introduced by Foldseek [27]. These distributions are analogous to sequence profiles derived from protein sequences compiled in multiple sequence alignments (MSAs) that describe protein families. While traditional MSAs proxy the diversity of sequence space of related proteins (so-called homologs), our description proxies the dynamic space of a single protein sequence within the constraints of the 3Di alphabet. We propose the term FlexProfiles to describe profiles of tokenized structure ensembles, which may be derived from both time-dependent and independent conformational data. Since the profiles employed here are derived from 3Di, we refer to these as 3Di-FlexProfiles. Our method expands the capabilities of pLMs by learning these coarse-grained protein-dynamics analogs through supervised, parameter-efficient Low Rank Adaptation (LoRA) fine-tuning [30]. By internalizing the coarse-grained information encoded in 3Di-FlexProfiles into the pLM’s model weights, our method bypasses the computational limitations of resolving high-resolution protein motions. Thereby, it captures coarse-grained protein dynamics directly from protein sequence in high-throughput and may be leveraged in large-scale workflows.

The resulting 3Di profiles reproduce observables from MD and improve remote homology detection compared to standard 3Di sequences.

### Summary of contributions

- We propose FlexProfiles as a coarse-grained representation of protein dynamics, and investigate how well 3Di captures conformational variability and dynamics.
- We introduce **ProtProfileMD**, the first framework that transfers MD-based 3Di profiles (FlexProfiles) into pLMs, enabling the prediction of protein dynamics proxies directly from amino acid sequences.
- We benchmark homology detection performance using Foldseek.

### Related work

The discretization of 3D structures into 1D strings over an alphabet by mapping backbone geometry to a structural alphabet has been used, in conjunction with substitution matrices, to enable fast structure alignment via string-based bioinformatics methods. These discretizations preserve useful structural and evolutionary signals [31, 27].

Foldseek introduced the 3Di alphabet, which describes tertiary residue-residue interactions and showed that converting structures to 3Di sequences enables orders-of-magnitude faster structure comparison and clustering with high sensitivity [27, 26]. Combining Foldseek with ProstT5, a pLM which directly translates between amino-acid and 3Di sequences, eliminates the need for experimentally determined or explicitly predicted structures [29].

PBxplore is the first method to provide a framework for translating conformations into protein blocks and computing per-residue frequency profiles and entropies across MD trajectories [32]. Previously, methods such as PETIMOT and SeaMoon used pLM embeddings to predict motion vectors indicative of dynamics from amino acid sequence [13, 33]. Such vectors help analyze functional motion in individual proteins. However, they do not enable large-scale searches, rendering those two solutions complementary to our new approach. The method most tangential to ours, ESMDynamic, trains ESMFold [34] to predict dynamics as C_*α*_ atom-derived contact probability maps, which function as global pairwise, probabilistic, time-averaged dynamic descriptors [35]. While capturing global flexibility, the pairwise nature of ESMDynamic prevents its use for large-scale comparisons of protein dynamics. Together, these methods substantiate the premise that MD-grounded targets, akin to our FlexProfiles, are learnable from sequence and pLM features.

## Materials and methods

### Profiles

Profiles provide per-residue statistical distributions that quantify how frequently each symbol in a given alphabet occurs at each position across a set of aligned or equal-length sequences. To avoid division by zero when converting raw counts into logodds, small background-probability-derived pseudo-counts may be added. Our method uses the common Position Probability Matrix (PPM). Let 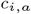 be the count of symbol *a* ∈ 𝒜, where 𝒜 is the alphabet, at alignment position *i*, with pseudo-count *α* derived from background frequencies, then an entry in the PPM is computed as follows:

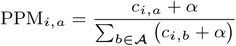

### Dataset

The simulations in *mdCATH* cover 5,398 protein domains [45], a subset of the *S20 homology set* of CATH release 4.2.0 [36], i.e., CATH clustered at 20% pairwise sequence identity (PIDE), which allows sampling known protein space more uniformly. Each protein domain in mdCATH is simulated with five replicas at five temperatures.

We computed the ground-truth FlexProfiles by discretizing each trajectory frame into a sequence over the 3Di structural alphabet using the Foldseek Vector Quantised-Variational AutoEncoder (VQ-VAE). Since all resulting 3Di sequences originated from the same protein, they are trivially aligned, enabling aggregation into 3Di-FlexProfiles that abide by the PPM definition (with *α* = 0). We performed this for all replicas and temperatures in mdCATH, treating duplicate sequences across temperatures and replicas as data augmentation.

We segmented the mdCATH dataset into disjoint training and evaluation partitions based on CATH version 4.2.0 structural hierarchies [36]. This ensured that the evaluation structures were not trivially similar to training structures. For this purpose, we selected the *Topology* hierarchy *h*, which should minimize structural overlap. Our dataset of size *N* = 5, 398 can be imagined as set of tuples 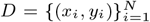, where *x*_*i*_ is a protein-domain and *y*_*i*_ is a CATH topology class label under *h*. We first compute our class sizes *t*(*c*) for the hierarchy class *c*.

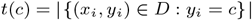

The selected classes 𝒮 chosen for the test set are obtained by excluding singletons, topologies with only one member, as well as very large classes (*>* 30 members), since these provide either no comparative signal or are overly dominant. From the remaining superfamilies, we sampled *k* = 50 at random without repeats to select the evaluation classes 𝒯 with a fixed random number generation seed *s* = 42.

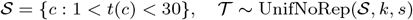

We created the evaluation dataset *D*_eval_ from *S*. All remaining entries were assigned to the training set *D*_train_.

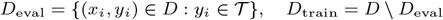

This procedure produced an evaluation dataset comprising approximately 3.56% of the data (192 entries), with the remaining 96.44% allocated to training (5206 entries). We further excluded domain *3rhaA00* from the train split, as it is a subdomain of *3rhaB03* in the test split (100% PIDE). The maximal PIDE between any protein in the training and testing set was 34.3%.

### Base model

ProtProfileMD builds on top of the ProstT5 pLM (**Pro**tein **St**ructure T5), which expands on the amino acid sequence-only ProtT5 [17] by introducing a bilingual formulation of protein learning wherein it translates amino acids to Foldseek’s 3Di structure tokens (”folding”) and vice versa (”reverse folding”) [29].

### Parameter-efficient finetuning

Instead of computing a full weight update, Low-Ranked Adaptation (LoRA) [30] constrains the update to the product of two low-rank decomposition matrices *BA* with rank *r* ≪ *d* and scaling factor *γ*, while the original weight matrix *W* remains frozen.

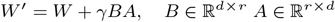

We enabled LoRA on attention projections with rank *r* = 8, *α* = 16, and dropout = 0.05, in the target modules *QV* (query and value projections) [30]. This optimization resulted in a total of 2,199,988 trainable parameters. We experimented with finetuning different parameter sets, e.g., *QKV O*, but did not observe meaningful improvements.

### Model architecture

ProtProfileMD (Figure 1) fine-tunes a pLM with a lightweight prediction head on top to predict, for each residue *i* in a sequence of length *L*, a row-stochastic probability distribution *P* ∈ ℝ^*L×*|𝒜|^, with | 𝒜| = 20 for the 3Di alphabet.

**Fig. 1:**
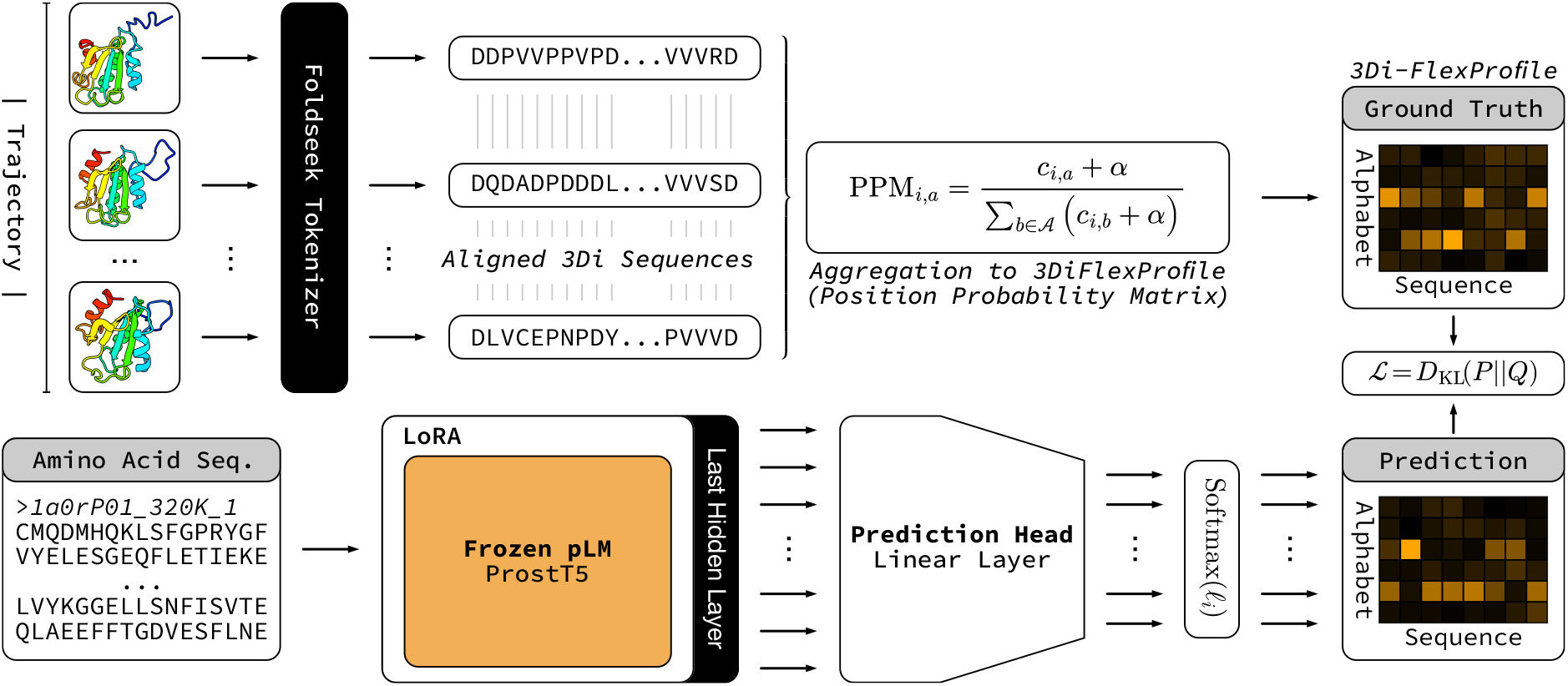
Schematic of ProtProfileMD data generation and training. ProtProfileMD learns residue-level state distributions directly from amino acid sequences by conditioning a pLM on 3Di-FlexProfiles, MD-derived 3Di sequence profiles. Distributions are obtained by discretizing MD trajectory frames from the mdCATH dataset into sequences over the discrete structural 3Di alphabet, introduced by Foldseek. The resulting sequences are subsequently aggregated into 3Di profiles as position probability matrices (PPMs). Amino acid sequences are encoded by passing the last hidden layer’s output to the ProstT5 pLM. The last hidden layer is passed to a prediction head, and an element-wise softmax is applied to produce a residue-wise probability distribution over the 3Di alphabet, resembling a 3Di profile. Training minimizes the KL divergence between predicted and ground-truth FlexProfiles. Gradient updates are applied to the pLM in a parameter-efficient manner via Low Rank Adaptation (LoRA).

**Fig. 2:**
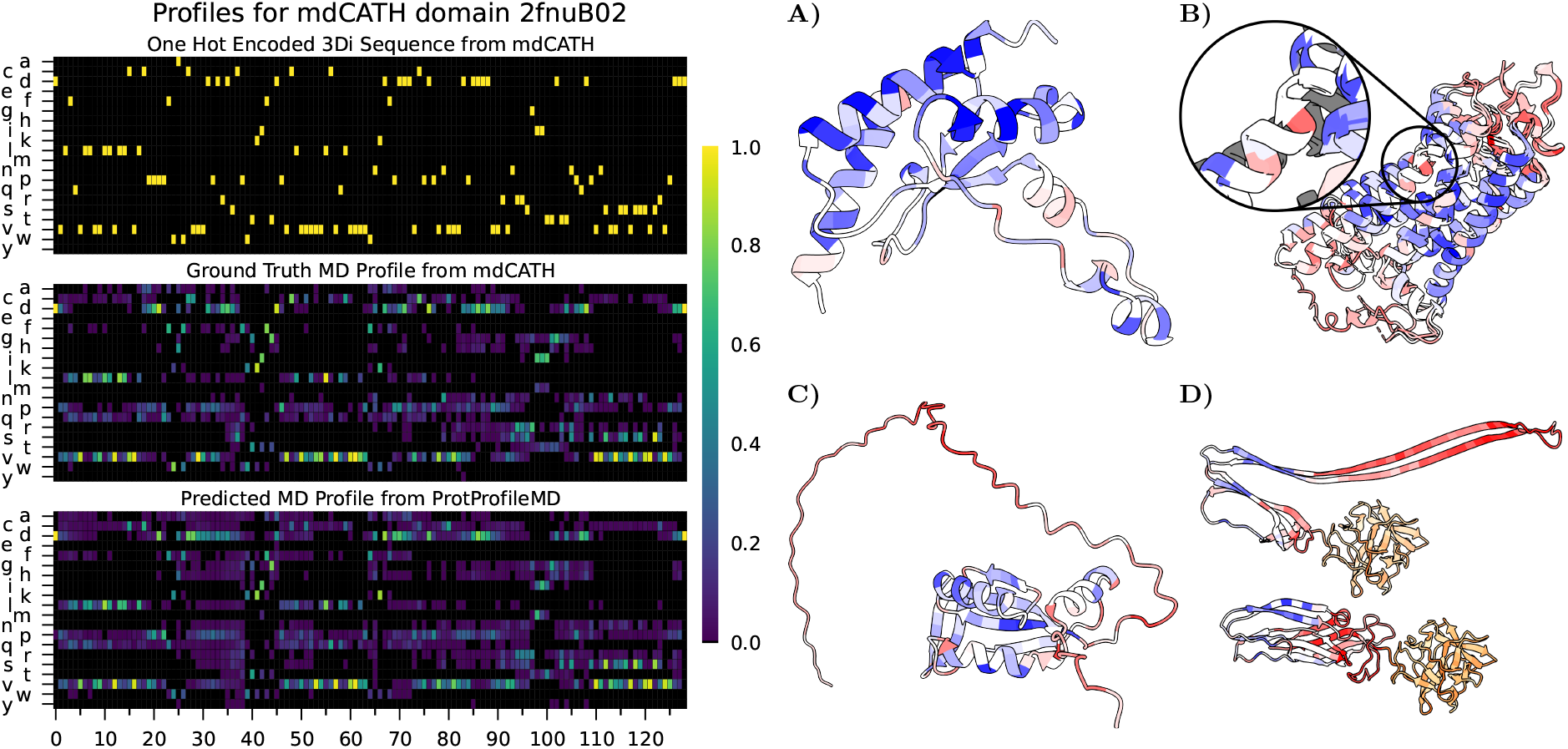
(Left) 3Di-FlexProfiles capture dynamic variability beyond discrete 3Di. The 3Di sequence from our evaluation split (CATH: *2fnuB02*) is represented as a static, one-hot-encoded profile, showing no variability (Top), while the ground-truth 3Di-FlexProfile (Center) and the predicted 3Di-FlexProfile (Bottom) have greater variability and higher information content. The color scale indicates probability values from 0 (black) to 1 (yellow), with columns (residue dimension) summing to 1. **(Right) Predicted 3Di-FlexProfiles distinguish functionally and dynamically different regions**. The color scale indicates per-residue entropy; high entropy regions are red, low entropy regions are blue. All panels (A–D) share a common color scale. (A) Stable/Rigid Protein Domain (CATH: *2fnuB02* [36]), from our mdCATH evaluation split, shows lower entropy in rigid regions as compared to more flexible regions. (B) G Protein-Coupled Receptor (GPCR) Bovine Rhodopsin, inactive and active states (PDB: *1F88* [37], *3PQR* [38]), shows high entropy in the region where conformation change occurs when activated (near highly conserved residue Proline 267 on helix 5 [39]). (C) Disordered Protein (AFDB: *O60888* [40]) shows high entropy in disordered regions, and low entropy in structured regions. (D) Fold switching proteins [41] are characterized by having largely the same amino acid sequence but significantly different structures deposited in the PDB [42]. Areas with large conformational divergence exhibit high entropy. Orange highlights the non-fold switching region (Mean entropy: 2.003*±*0.474 bits), and blue to red (PDB: *5EC5 [43]* (Top), *3ZXG* [44] (Bottom)) annotate the fold switching region (Mean entropy: 3.065*±*0.224 bits).

We first tokenized, then embedded the input amino-acid sequences by extracting the last hidden layer *H* ∈ ℝ^*L×d*^, *d* = 1024 from the pLM encoder, *ProstT5*. The encoder model weights *W* were updated with LoRA. A linear transformation mapped *H* to class logits *ℓ* ∈ ℝ^*L×A*^. We experimented with more complex prediction heads but did not observe meaningful improvements, so we used the simplest models in line with Occam’s razor. A dropout of 0.1 was used for the prediction heads during training on the last hidden layer.

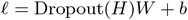

We applied the softmax function to each residue dimension to constrain the probability distribution over possible structural states (i.e., the class dimension).

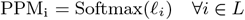

The predicted FlexProfile can be used in downstream pipelines without further postprocessing.

### Batching, masking, and padding

Batches were sampled randomly, without replacement, from the training set *D*_train_. We set the attention mask to *M* ∈ {0, 1}^*L*^. Each batch was padded to the longest sequence in the batch to account for non-equal-length input sequences and extended to a multiple of 8 to increase computational efficiency on modern hardware. The Hadamard product zeros out masked positions in the raw embeddings before passing *H* to the prediction head, yielding *H*^*′*^ = *M* ⊙ *H*, where *H*^*′*^ ∈ ℝ^*L×d*^ contains only valid token representations. The ProstT5 special token vector *h*_*i*=0_ ∈ *H*^*′*^ was removed for all sequences in a batch.

### Learning objective

Training minimizes the Kullback-Leibler (KL) divergence *D*_KL_(*P* ∥Q) [46] between the target per-residue distribution *P* and the predicted per-residue distribution *Q*, where *L* is the number of valid residue positions.

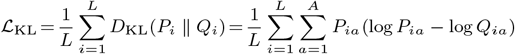

After masking out non-token positions, we flattened the batch across the residue dimension, such that the KL-divergence was weighted uniformly across all residues. We passed *P* and log *Q* directly to PyTorch’s *KLDivLoss* with the batchmean reduction.

### Training

We trained our final model for five epochs, totaling approximately three A100 GPU hours across all replicas at the lowest temperature (320 Kelvin). We used a per-device batch size of five across four devices running in distributed data parallelism, with ten gradient accumulation steps, yielding an effective batch size of 200. We used *AdamW* [47], with default PyTorch hyperparameters and an initial learning rate of *η*_0_ = 10^™3^. A linear warm-up of 200 steps was followed by a cosine decay scheduler to smoothly anneal the learning rate to zero. We logged step-wise metrics and tracked runs in Weights & Biases [48]. Checkpoints were saved every 300 steps, and the best (lowest KL-divergence on the validation set) was selected at the end.

### Evaluation

To assess whether 3Di discretization of MD simulations captured global conformational variability and drift, we compared 3Di substitution-weighted Hamming distances with root-mean-square deviation (RMSD) and FlexProfile entropy [49] with root-mean-square fluctuation (RMSF). RMSD measures the continuous Euclidean displacement of atoms after rigid-body alignment, whereas Hamming distance quantifies sequence-level divergence in the 3Di alphabet. By weighting substitutions according to the 3Di substitution matrix, we ensured that similar local 3Di states were penalized less severely than dissimilar ones. RMSF quantifies positional variability of atoms across a trajectory, whereas entropy reflects the variability of 3Di state distribution sampled at each residue. RMSD and Weighted Hamming distances were computed per frame with respect to the first frame and averaged across trajectories. Predicted 3Di-FlexProfiles were compared with ground-truth at 320K, the same temperature used to train the model. Entries for the reference frame were omitted, since these do not assume meaningful values for RMSD and hamming distance.

We used UCSF ChimeraX [50] to visualize how the per-residue entropy of FlexProfiles corresponds with 3D structures for various protein classes of interest.

## Results

We showed that Foldseek’s [27] discrete structural alphabet, 3Di, can capture conformational changes of proteins derived from MD simulations, and that salient evolutionary information may be contained in and extracted from protein dynamics. Analogous to conventional protein sequence profiles, which capture amino acid frequencies over homologs in a protein family, we propose *FlexProfiles*, which capture discrete structure token frequencies over conformational ensembles of MD trajectories of a protein. As we use Foldseek’s 3Di alphabet here as discrete 3D descriptors, we term the corresponding profiles *3Di-FlexProfiles*. Our specific implementation for deriving such profiles, ProtProfileMD, bypasses the need for computationally expensive MD simulations by predicting FlexProfiles directly from single protein sequences, leveraging knowledge encoded in pLMs (Figure 1).

When used for remote homology detection, we observed increased sensitivity with FlexProfiles when combined with current structure comparison tools, which, to date, could not incorporate dynamics.

### FlexProfiles capture protein dynamics

Using Foldseek’s 3Di alphabet to represent each frame and protein in mdCATH [45], we demonstrated that coarse-grained structure tokenizers retain sufficient information to capture some aspects of protein dynamics. By correlating the root-mean-square-deviation (RMSD) of each MD frame with respect to the first frame (Section 2.9) against the substitution-weighted Hamming distance, 3Di correlated with conformational changes (Pearson: 0.65 to 0.79, depending on simulation temperature, Fig. 3A). The 3Di also captured local residue-level fluctuations because the average flexibility of each residue (measured as RMSF, Section 2.9) correlated with the Shannon entropy of the per-residue 3Di-FlexProfile frequencies (Pearson: 0.76 to 0.87, Fig. 3B).

**Fig. 3:**
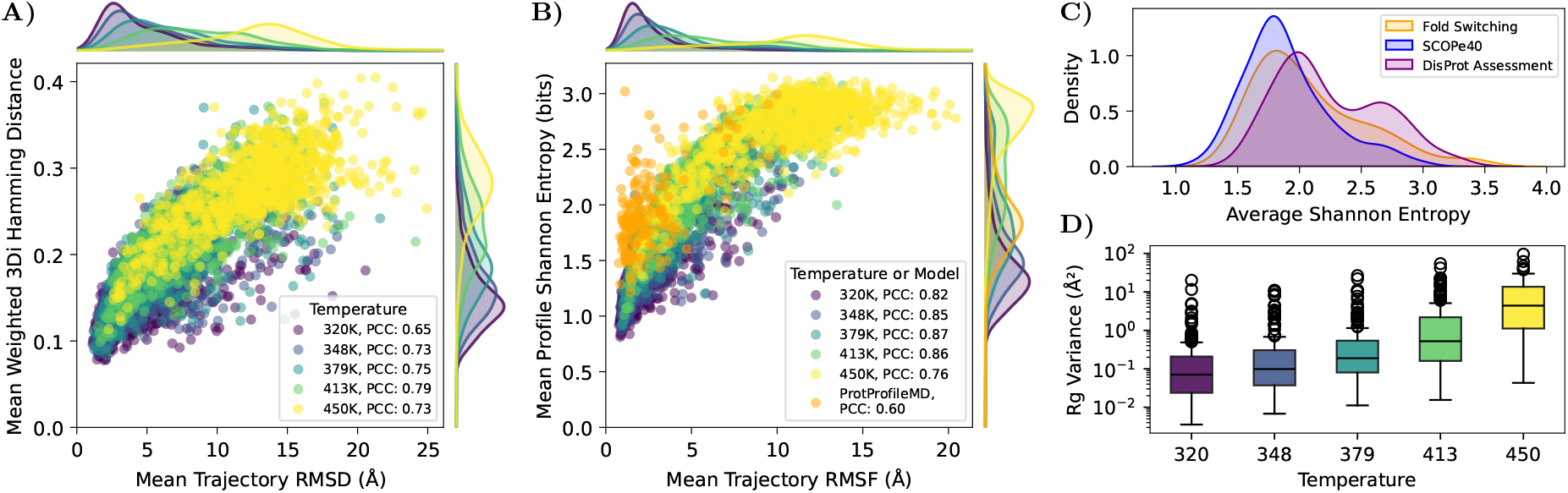
Information-theoretic signals from 3Di profiles, recapitulate MD-derived structural dynamics observables. (A) Averaged per-frame RMSD w.r.t. the first frame correlates well with the 3Di substitution matrix weighted hamming distance of each frame’s 3Di sequence w.r.t. the first frame, for 3Di sequences computed directly from simulation data of our evaluation split. (B) Per-trajectory-averaged RMSF correlates well with the averaged per-residue profile entropy for profiles inferred from simulations. Predicted profiles correlated with the RMSF from simulations at 320K, albeit at a lower level as compared to groundtruths (Orange). (C) The average Shannon entropy of predicted 3Di-FlexProfiles increased when comparing protein classes with increasing flexibility as proxied by comparatively rigid (SCOPe40 [51, 52]), in-between (Fold Switching [41]), and flexible proteins (DisProt Assessment [40]). (D) The observed variance of the radius of gyration between replicas in the ground truth MD simulations increased with increasing simulation temperatures, highlighting growing deviations between replicas at higher temperatures. This informed the decision to use data exclusively from the lowest temperature (320K) for training ProtProfileMD.

### Sequence alone predicts 3Di-profiles

We investigated whether or not sequence information alone suffices to predict 3Di-Profiles by finetuning an existing pLM (ProstT5). Towards this end, we used 3Di-FlexProfiles derived from mdCATH (Figure 1, and Section 2.1) to parameter-efficient (LoRA) finetune ProstT5 to directly predict for each residue in a protein a distribution over 3Di states, inputting amino acid sequences.

We selected to train our model only on profiles generated from the lowest simulation temperature of 320K. This decision was informed by an increasing variance in the radius of gyration at higher temperatures, suggesting greater deviations from the folded ensemble and increased global expansion (Figure 3D). Additionally, unfolding trajectories became more common at higher temperatures, which further decreased the signal-to-noise ratio. Higher temperatures inflate entropy by injecting unstructured variance that the model cannot meaningfully infer from a sequence.

On a strict hold-out set created by splitting the mdCATH data into CATH-derived topology-splits (2.2), we observed a good agreement between the Shannon entropy of MD-derived 3Di distributions and RMSF (Pearson=0.82 for the evaluation split at the temperature selected for training, Figure 3).

The distribution of average Shannon entropy of predicted 3Di-FlexProfiles shifts upward with increasing protein flexibility. Rigid SCOPe40 domains concentrate at lower entropies, while fold-switching and DisProt proteins show broader, higher-entropy distributions (Figure 3C).

### Predicted 3D-Profiles boost remote homology detection

Leveraging ProtProfileMD, we applied predicted FlexProfiles to improve the detection of remote homologs (protein pairs with similar structure and dissimilar sequence), without explicit structure prediction. For this, we replicated Foldseek’s benchmark across four different modalities: (1) amino acid sequence search with MMseqs2 [28], and 3Di search with Foldseek [27] using either (2) 3Di derived from PDB structures, or (3) 3Di predicted using ProstT5, and (4) finally the 3Di-FlexProfile search introduced here using predictions from ProtProfileMD. The ROC-AUC as a function of sensitivity up to the first false-positive, plotted against the fraction of queries, shows that our new method, rooted in dynamics, improved across all SCOPe levels, i.e., *Family, Superfamily*, and *Fold* (Figure 4).

**Fig. 4:**
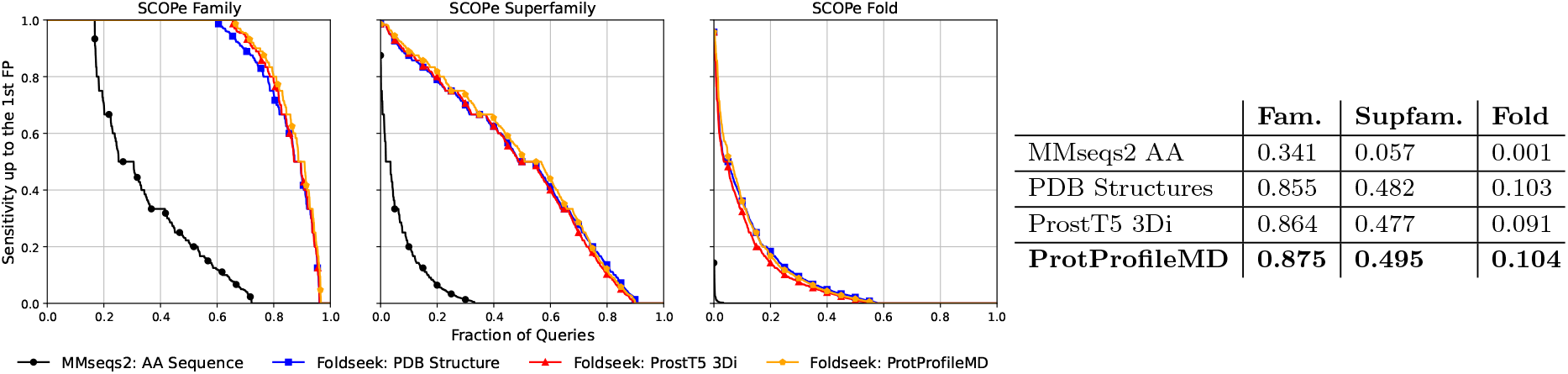
Searching with FlexProfiles improves remote homology detection as measured by the SCOPe Sensitivity Benchmark. Following the Foldseek benchmark, we plot the cumulative sensitivity up to the first false positive against the fraction of queries covered when searching SCOPe against itself using either MMseqs2 (amino acids), or Foldseek (3Di derived from PDB structures, 3Di predicted using ProstT5, or the 3Di-FlexProfiles predicted with the proposed ProtProfileMD). Performance is shown for the three hierarchical levels of the SCOPe classification: Family (left), Superfamily (center), and Fold (right). Detailed performance values are given in the table on the right.

## Discussion

A mechanistic understanding of how amino acid sequences give rise to both structure and conformational dynamics, and where dynamic patterns are evolutionarily conserved, is increasingly central to bridging low- and high-throughput experiments using homology detection and broadening our understanding of the sequence-structure-function paradigm. Scalable protein structure comparison [27], enabled by high-resolution 3D structure predictions [53], has markedly expanded our ability to detect associations previously deemed undetectable. Although proteins populate conformational ensembles and dynamics are often integral to function, most ubiquitous structure-comparison methods still dismiss this additional level of complexity and operate on single static representations, typically one PDB or AlphaFold model per protein.

Here, we investigated whether incorporating a simplified aspect of conformational dynamics can enhance homology detection by expanding discrete structure tokens to probabilistic representations. Instead of exclusively encoding a single dominant and stable state, we derived per-residue probabilities over the 3Di states introduced by Foldseek [27], from conformational changes observed during MD simulations. We dubbed these distributions *FlexProfiles*. Unlike conventional sequence profiles, which are family-averaged as they capture sequence or structural plasticity between related proteins, the proposed FlexProfiles capture protein dynamics derived from a single protein, and are consequently protein-specific. Despite discarding atomic-level detail, our specific implementation, using Foldseek’s 3Di alphabet-based *3Di-FlexProfiles*, preserves residue-level conformational variability, highlighting rigid (low entropy) and flexible (high entropy) regions. 3Di-FlexProfiles provide a detailed, biologically and biophysically faithful representation of the conformational landscapes, consistent with established MD observables, such as RMSF (Pearson: 0.6, Figure 3B).

Simulating proteome-sized datasets to derive FlexProfiles remains computationally intractable, and emerging generative models that seek to bypass this restriction, e.g., by sampling ensembles, are still not deployable at scale. Analogously, we showed that structure prediction is not a prerequisite for leveraging structure comparison sensitivity by directly translating amino acid sequences into 3Di sequences [29, 26, 54]. Following this logic, we finetuned the same protein language model (ProstT5 [29]) to map amino acid sequences directly to the proposed 3Di-FlexProfile with conformations derived from MD simulations.

Replicating Foldseek’s remote homology detection benchmark, using ProtProfileMD to predict 3Di-FlexProfiles has improved sensitivity compared to searching with native 3Di. Although the improvements are consistent, the gain remains marginal compared to searching with 3Di-sequences. Nonetheless, this advance comes with little to no additional cost, as throughput is virtually identical to the original ProstT5 (which required around 35m, or 0.1s/protein, to process the entire human proteome on a single RTX A6000 GPU).

Our novel approach is not limited to a particular methodology for measuring protein flexibility, structural alphabet, or model variant. Instead, it defines a proof-of-principle, generic strategy for coupling probabilistic summaries from physical simulations or other modalities with discrete structural alphabets and pLMs. Our results support the view that FlexProfiles and the prediction thereof can enable novel applications, such as proteome-scale annotation and search for proteins or motifs without rigid structure.

Our *ad hoc* solution seems to leave ample room for improvements. These might include the following: a) expand training data (mdCATH consists only of single domains, and MD-derived motions are limited in spatial and temporal scale), b) apply more demanding benchmarks (SCOPe contains comparatively rigid domains), c) refine structural alphabets (20 3Di-states could be insufficient to capture the full dynamic variability), d) advance search implementations (Foldseek does not yet support profile-profile comparison [27] - we performed profile-consensus search (Section 2.6)).

Dynamics-aware profiles could increase the sensitivity of remote homology detection and facilitate hypothesis generation about protein motion in the conformational landscapes that proteins explore in nature. Holistically, these results argue for a shift in the discussion: instead of prompting a model for a static protein structure, a researcher may inquire how structural states are distributed across residues, and then use that distribution to search, compare, learn, and design.

## Acknowledgments

We thank the mdCATH team for making their molecular dynamics simulations publicly available, which was essential for this work, and the CATH team for decades of dedication in categorizing protein domains. Thanks to David Wagemann and George Bouras for great discussions, Tobias Olenyi for technical support, and to Nikita Kugut (TUM) for support with many other aspects of this work. Last, but not least, we would like to thank all computational colleagues who make their tools available, all experimental colleagues who help advance science by making their data publicly available, and all those who maintain the resulting databases.

## Data availability

Code is available on GitHub at https://github.com/finnlueth/protprofilemd. Models and data can be found on HuggingFace at https://huggingface.co/finnlueth/protprofilemd and https://huggingface.co/datasets/finnlueth/protprofilemd.

## Author contributions statement

F.L. and M.H. conceived the experiments, F.L. conducted the experiments, and F.L. and M.H. analyzed the results. F.L. wrote the manuscript. M.H. edited the manuscript. M.H., B.R., M.M., and M.S. reviewed the manuscript. V.M. helped with Foldseek profile conversion. M.M. helped with Foldseek SCOPe by enabling the parsing of profiles and 3Di FASTA files to Foldseek databases. M.S. contributed valuable insights about profiles, probabilities, and Foldseek.

